## Supplemental Materials for "Climate predicts geographic and temporal variation in mosquito-borne disease dynamics on two continents"

**Table S1: Effect of temperature on dengue transmission dynamics from prior studies.** This table provides information used in Figure 5 of the main text to visualize the relationship between temperature and dengue risk compared with the relative basic reproductive number derived from the trait-based model. Asterisks indicate that the mean temperature was calculated as the average value of the minimum and maximum temperatures provided in the text (measured or estimated from temperature plots). Coefficient values are the values calculated in regression models in the sourced references, they indicate the estimated effect on dengue cases given a one-unit change in the temperature metric.

| Study location | Dengue metric | Mean temperature | Temperature metric | Temperature metric time lag | Coefficient value | Source |
|---|---|---|---|---|---|---|
| Mexico | Weekly cases | 14* | Weekly minimum | None | 0.44 | [1] |
| Mexico | Weekly cases | 15* | Weekly minimum | None | 0.58 | [1] |
| Mexico | Monthly cases | 16.5* | Monthly minimum | 4-8 weeks | 0.03-0.15 | [2] |
| China | Monthly cases | 19.5 | Minimum | 4 weeks | 0.732 | [3] |
| Guadeloupe | Yearly or seasonal cases | 22* | Minimum | 5 weeks | 0.108 | [4] |
| Vietnam | Monthly cases | 22.5* | Monthly mean | 8 weeks | 0.23 | [5] |
| Thailand | Monthly cases | 23 | Monthly minimum | None | 0.99 | [6] |
| Bangladesh | Monthly cases | 23* | Mean | 4 weeks | 6.07 | [7] |
| Mexico | Weekly incidence per 100,000 people | 25 | Mean weekly Sea Surface Temperature | 5 weeks | 0.2 | [8] |
| Guadeloupe | Yearly or seasonal cases | 26.5* | Mean | 11 weeks | 0.228 | [4] |
| Taiwan | Monthly cases | 29* | Deviation between monthly maximum and mean | None | -0.126 | [9] |
| Bangladesh | Monthly cases | 30* | Monthly maximum | None | 0.0105 | [10] |
| Vietnam | Monthly cases | 33.5* | Monthly mean | 8 weeks | -1.94 | [5] |
| Colombia | Monthly or Epi-period cases | 32 | Monthly mean | None | 0.001 | [11] |
| Thailand | Monthly cases | 35* | Monthly maximum | 4 weeks | -0.609 | [12] |

**Table S2: Number of days with interpolated temperature, rainfall, and humidity by site within study period.** There were 1,918 days in total for sites within Ecuador and 2,069 days in total for sites within Kenya. Interpolated rainfall values indicate days where there was one or more missing days with a rainfall value in the prior 14 days (inclusive of measurement date).

| Country | Site | Days with interpolated temperature | Days with interpolated monthly rainfall | Days with interpolated humidity |
|---|---|---|---|---|
| Ecuador | Huaquillas | 331 | 443 | 331 |
| Ecuador | Machala | 6 | 78 | 306 |
| Ecuador | Portovelo | 490 | 601 | 489 |
| Ecuador | Zaruma | 974 | 434 | 322 |
| Kenya | Chulaimbo | 289 | 511 | 288 |
| Kenya | Kisumu | 325 | 506 | 305 |
| Kenya | Msambweni | 142 | 602 | 142 |
| Kenya | Ukunda | 443 | 623 | 867 |

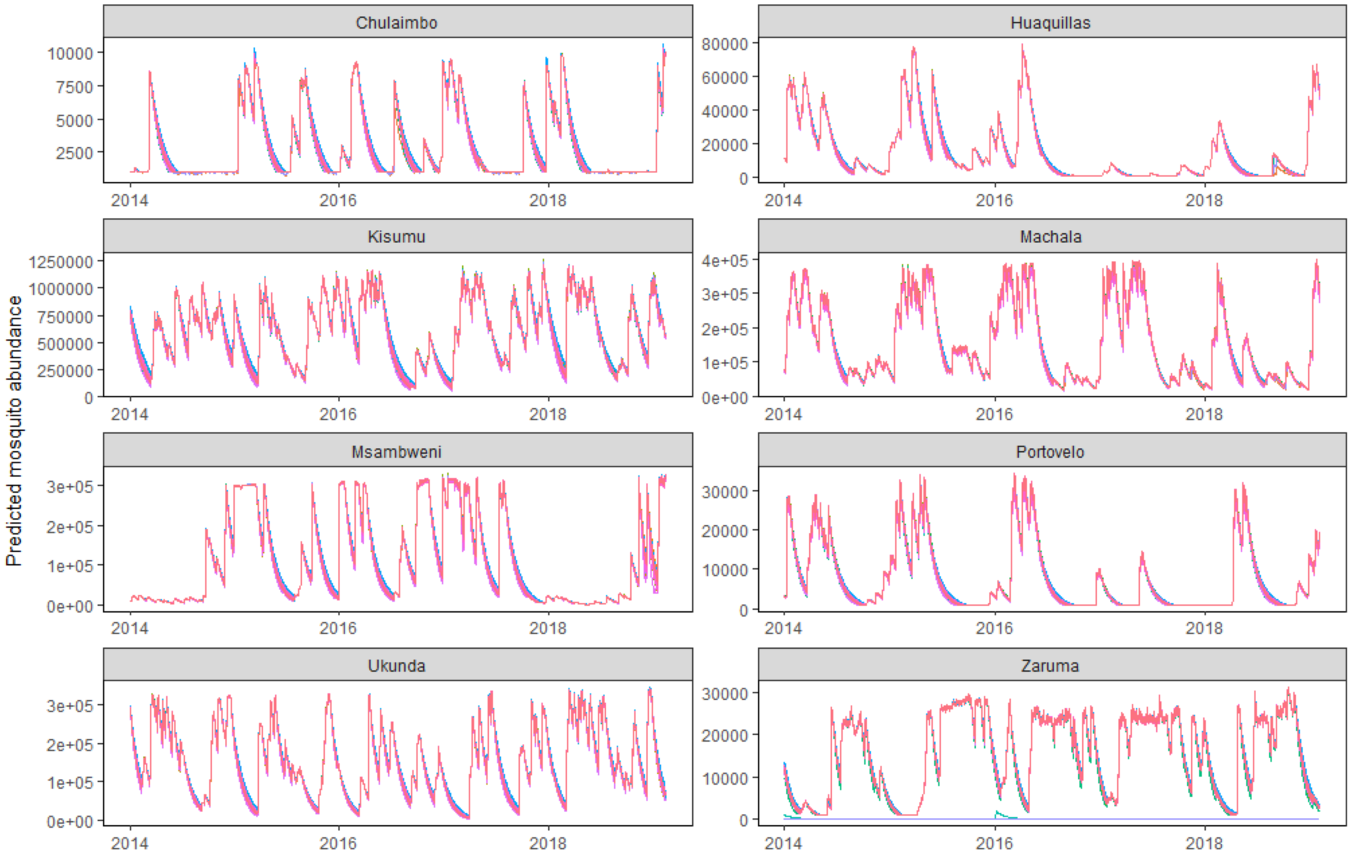


**Figure S1: Mosquito predictions are not affected by critical thermal minimum, maximum, and rate constants used in *Aedes aegypti* life history traits.** To determine the extent to which variations in *Aedes aegypti* life history traits affect model predictions, we conducted a sensitivity analysis. For the sensitivity analysis, we randomly sampled 50 different c, $T_0$, and $T_m$ estimates for temperature-dependent traits from the posterior distributions found in [13] and ran the SEI-SEIR model with those estimates. This plot shows the total predicted mosquitoes ($N_m$) for each site through time. The results for each model run are shown as a different colored line.

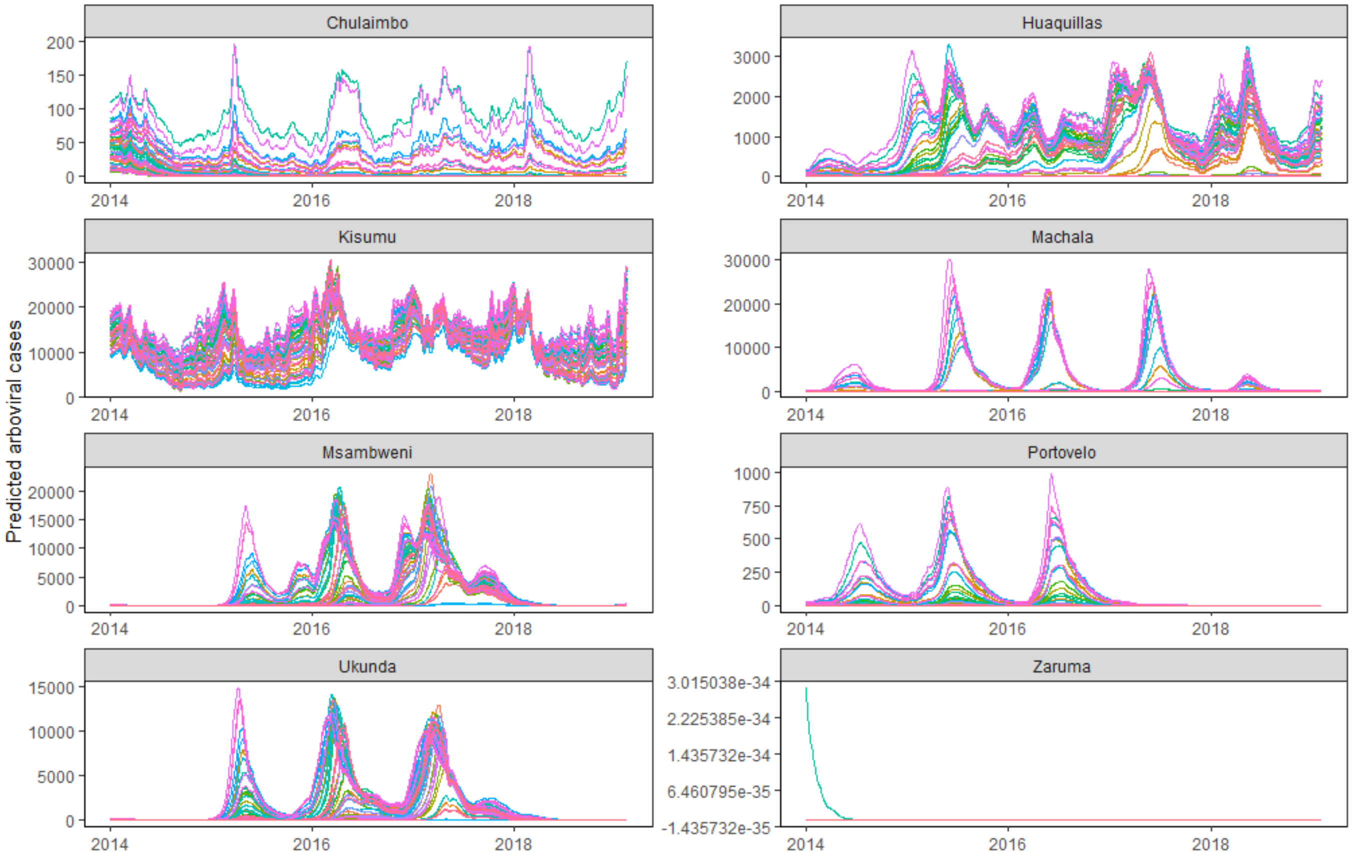


**Figure S2: Trajectories of predicted arboviral cases are not affected by critical thermal minimum, maximum, and rate constants used in *Aedes aegypti* life history traits.** To determine the extent to which variations in *Aedes aegypti* life history traits affected model predictions, we conducted a sensitivity analysis. For the sensitivity analysis, we randomly sampled 50 different c, $T_0$, and $T_m$ estimates for temperature-dependent traits from the posterior distributions found in [13] and ran the SEI-SEIR model with those estimates (same models as shown in Figure S1). This plot shows predicted cases ($I_h$) for each site through time. The results for each model run are shown as a different colored line. The different trait values varied the magnitude of predicted cases in many settings, but not the temporal dynamics. Many simulations overlap for Portovelo and Zaruma.

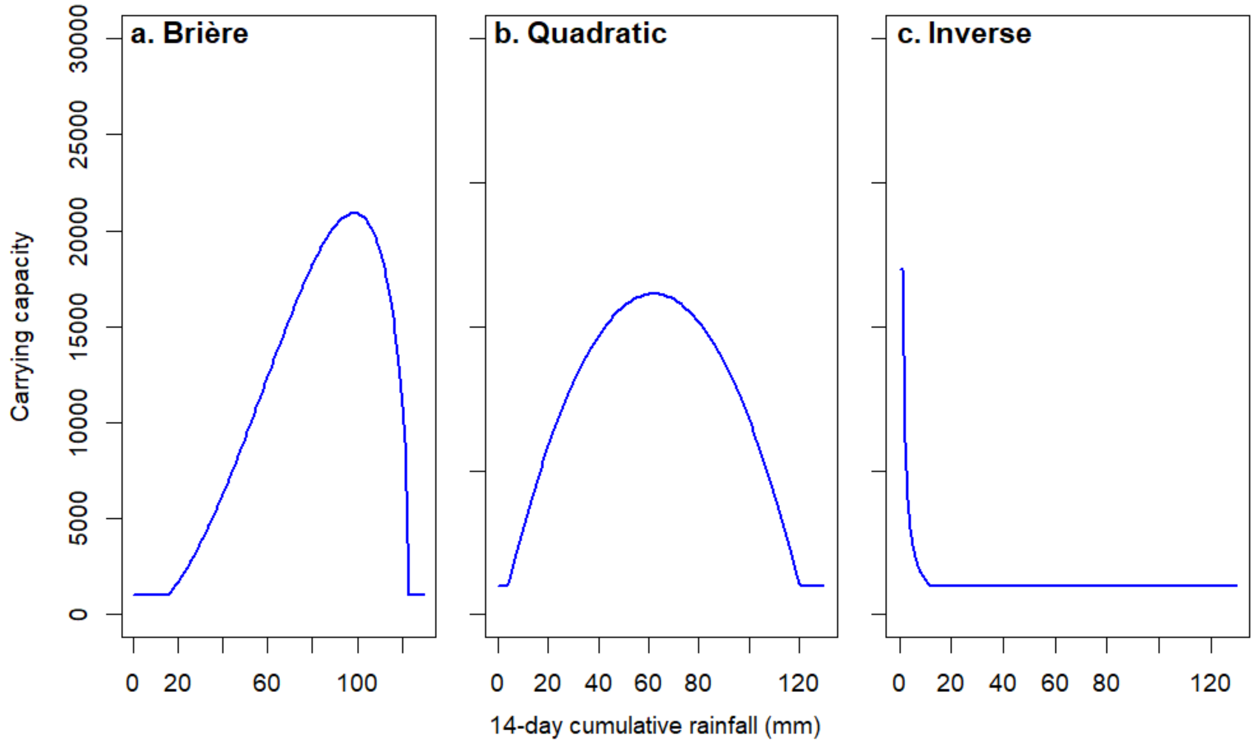


**Figure S3: Hypothesized functional forms for effects of rainfall on mosquito carrying capacity.** We tested three hypothesized functional relationships between 14-day cumulative rainfall values (following [14]) and mosquito carrying capacity: (a) Brière, in which carrying capacity increases with increasing rainfall until a threshold where flushing occurs; (b) quadratic, in which carrying capacity peaks at intermediate rainfall values, similar to [15]; and (c) inverse, in which mosquito abundance is greatest during periods of drought, similar to [14]. In these plots, temperature, maximum rainfall, and the human population are held constant at 29°C, 123 mm, and 20,000 people, respectively. See Methods for functional form equations.

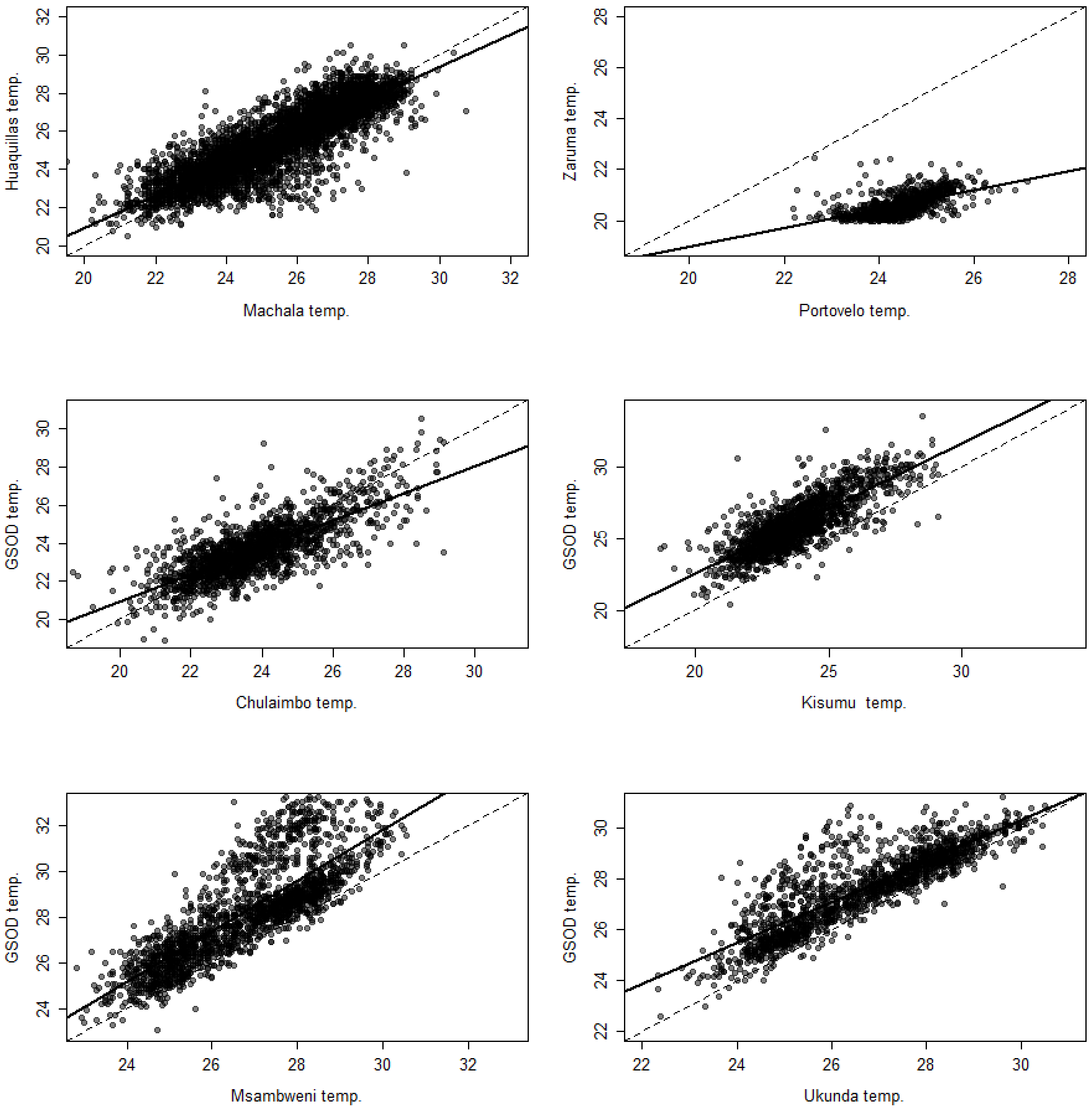


**Figure S4: Scatterplot comparing ambient temperatures measured at study sites with datasets used for interpolation.** For Ecuador, we used the nearest study site values when possible or else the long term mean temperature values for the corresponding Julien day. For Kenya, we used NOAA Global Surface Summary of the Day datasets from Kisumu Airport for Kisumu and Chulaimbo and from Mombasa Airport for Msambweni and Ukunda. Dashed lines indicate the y = x line and solid lines indicate the linear regression line used to interpolate data.

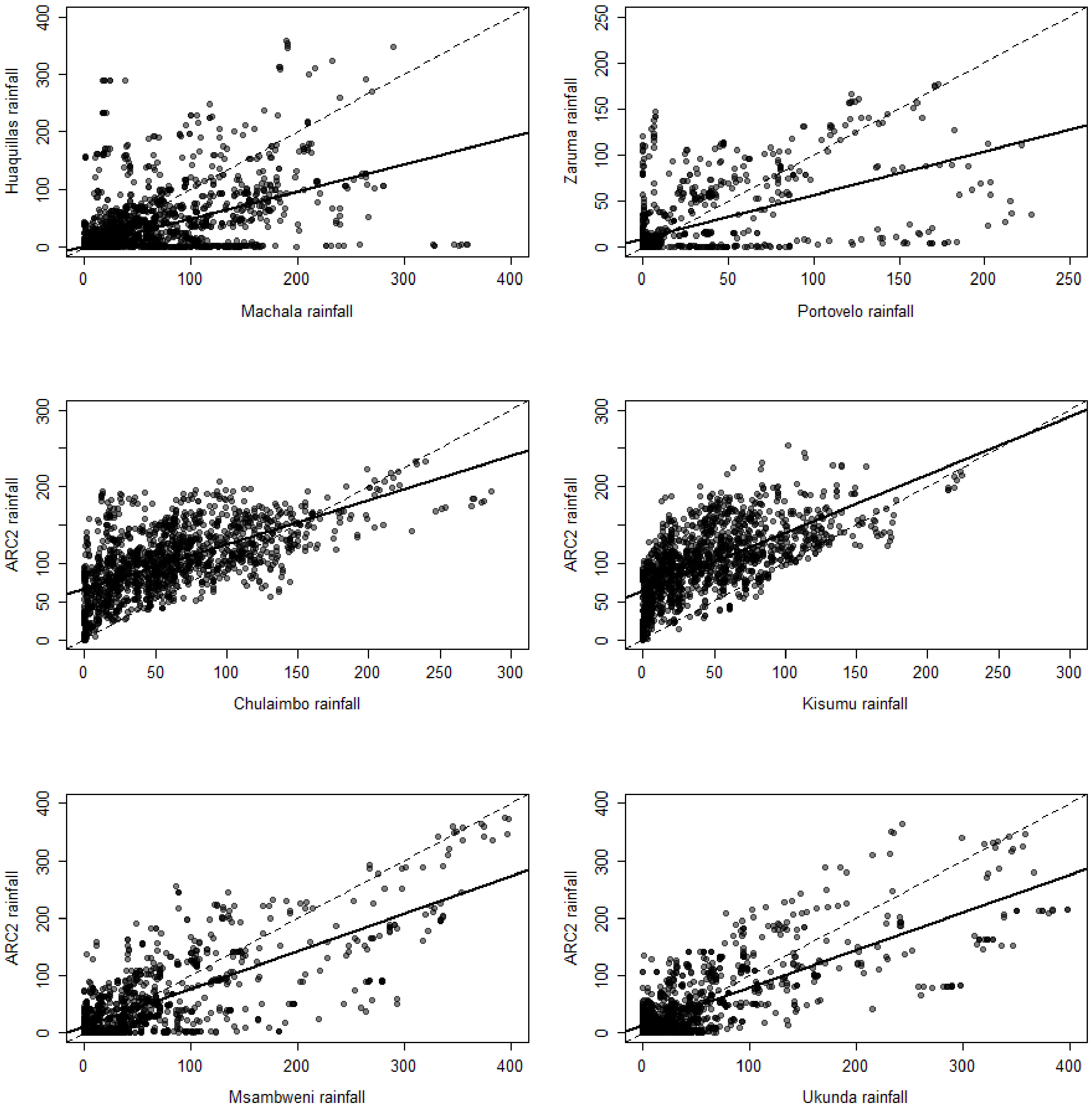


**Figure S5: Scatterplot comparing daily values for cumulative rainfall in the prior 14 days measured at study sites with datasets used for interpolation.** For Ecuador, we used the nearest study site values when possible or else the long term mean 14-day cumulative rainfall values for the corresponding Julien day. For Kenya, we used NOAA Global Surface Summary of the Day datasets from Kisumu Airport for Kisumu and Chulaimbo and from Mombasa Airport for Msambweni and Ukunda. Dashed lines indicate the y = x line and solid lines indicate the linear regression line used to interpolate data.

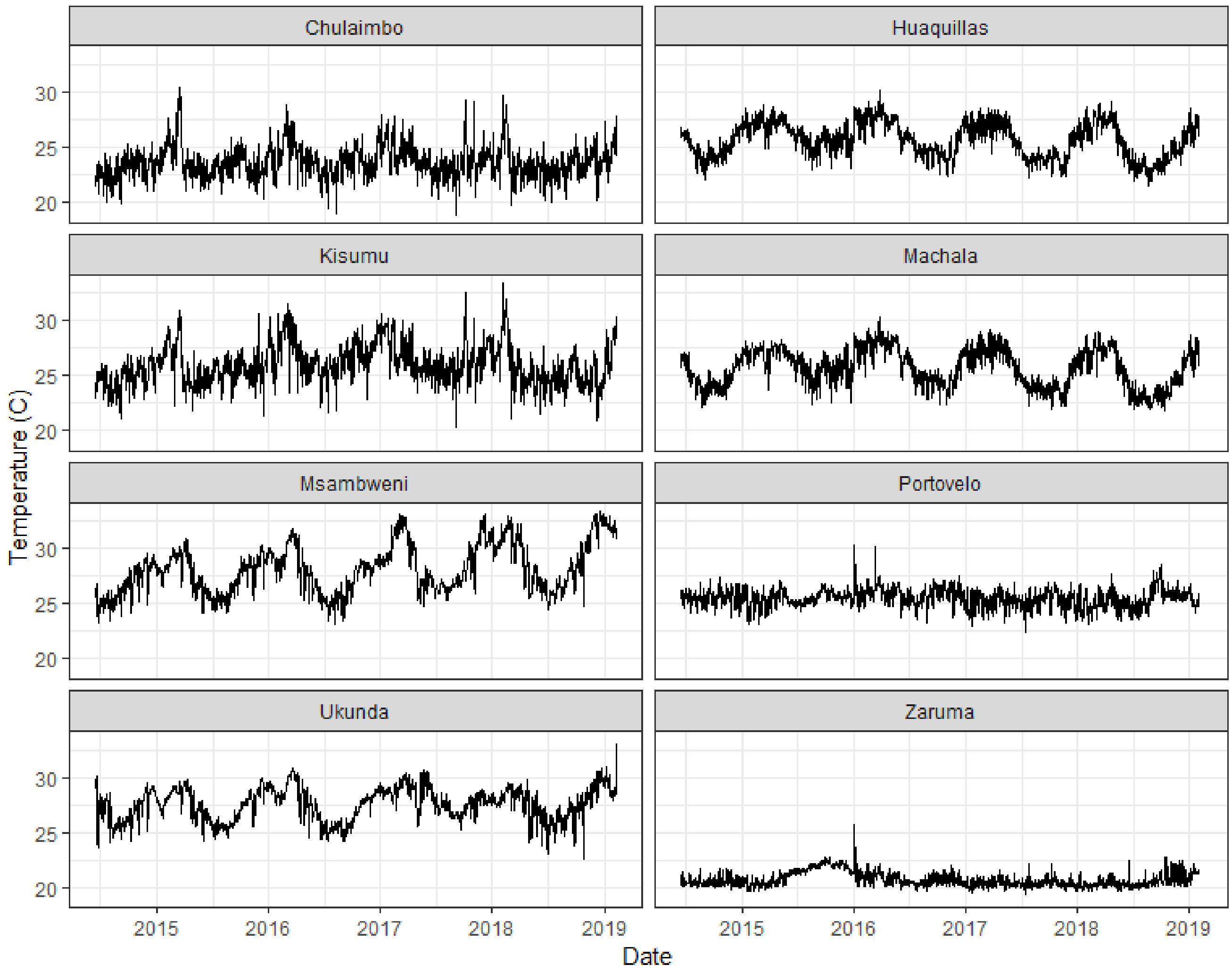


**Figure S6: Daily temperatures across sites within study period.** Kenya sites are on the left and Ecuador sites are on the right.

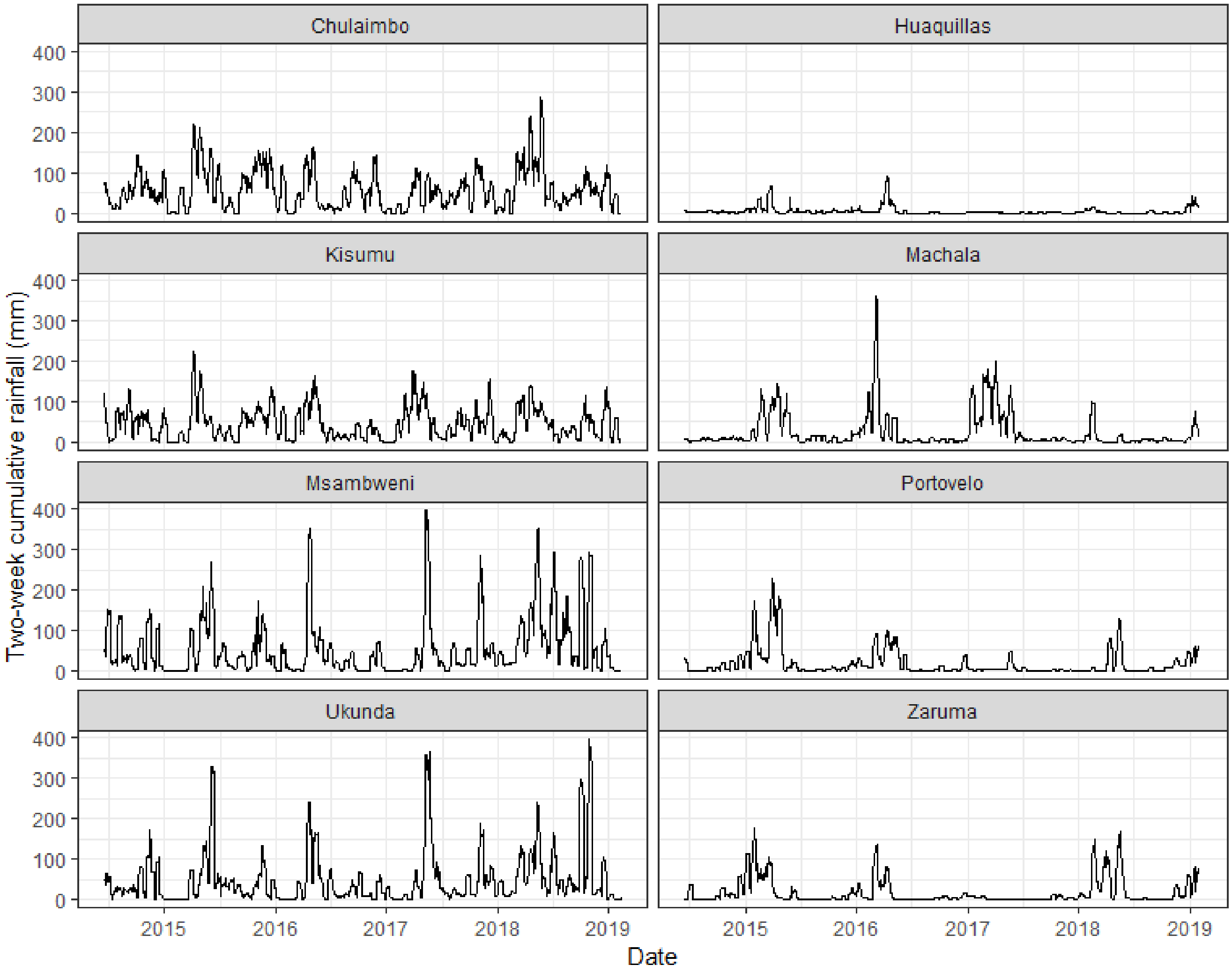


**Figure S7: Daily values of cumulative 14-day rainfall across sites within study period.** Kenya sites are on the left and Ecuador sites are on the right.

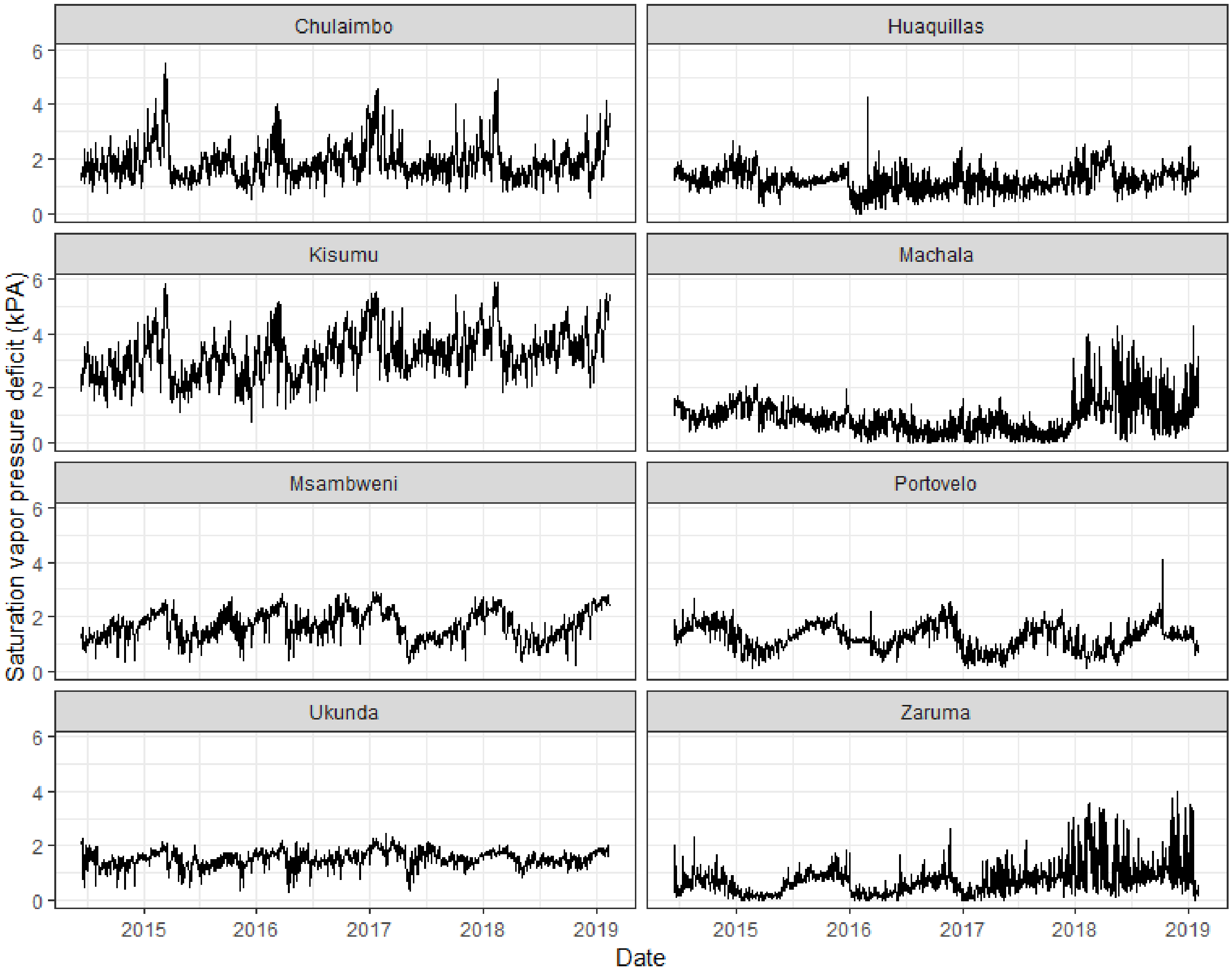


**Figure S8: Daily saturation vapor pressure deficit across sites within study period.** Kenya sites are on the left and Ecuador sites are on the right.

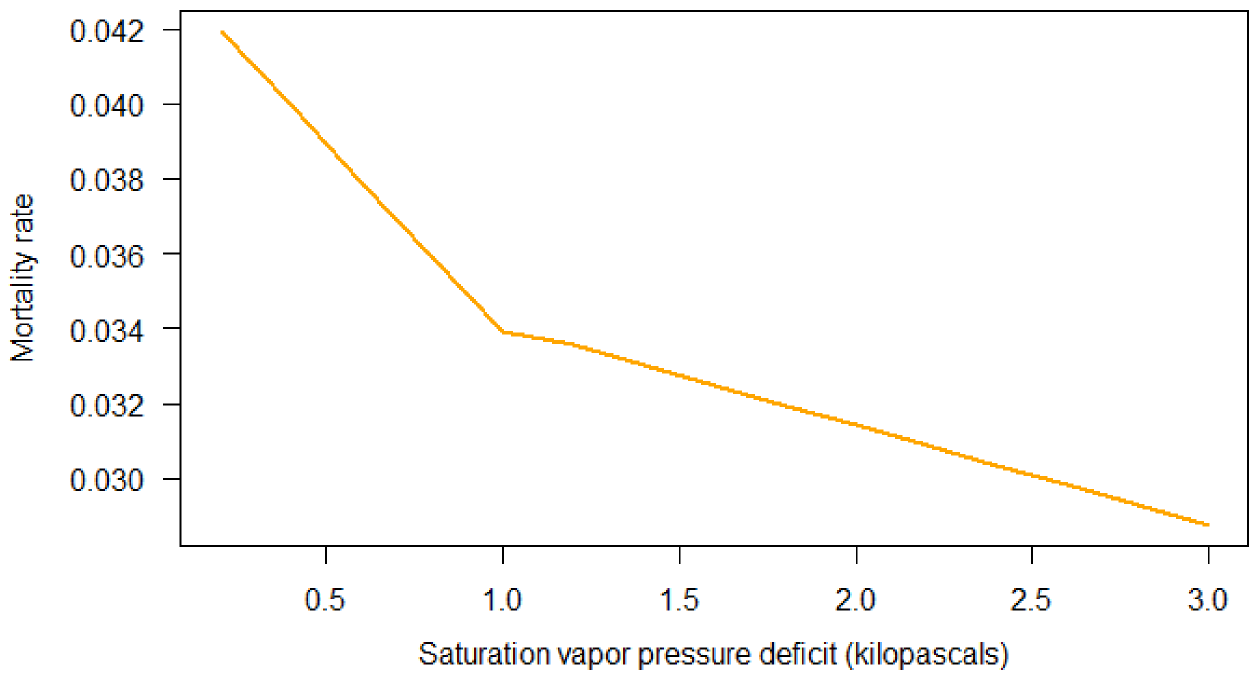


**Figure S9: Mosquito mortality rate as a function of saturation vapor pressure deficit (SVPD).** This relationship is a step function where the slope of the linear relationship is steeper for SVPD ≤ 1 compared with SVPD > 1. The step function is also scaled differently for SVPD ≤ 1 and SVPD > 1 to restrict the mortality rate within the range of mortality rates observed in other studies; these scaling factors make the function appear nonlinear between 1.0 and 1.2 in the plot.

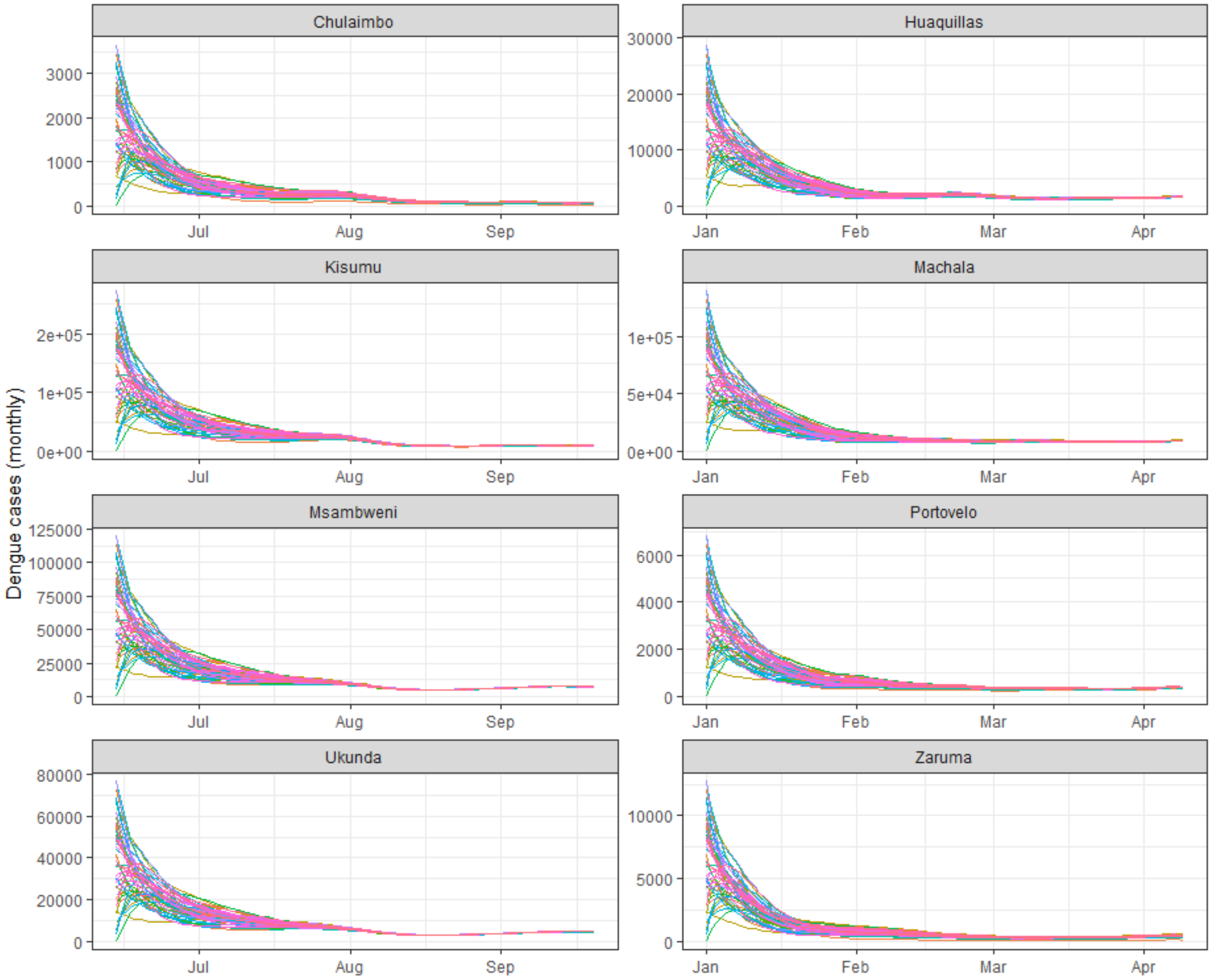


**Figure S10: Models converge after ~90 days regardless of initial conditions.** To determine how the model's initial conditions affected the magnitude and trajectories of each compartment and across sites, we conducted a sensitivity analysis. We randomly sampled 50 different proportions of starting conditions for each compartment where $S_V + E_V + I_V = 1$ and $S_H + E_H + I_H + R_H = 1$ using Latin hypercube sampling. Latin hypercube sampling is a statistical method for generating a random sample of values from a multidimensional distribution. We used the optimumLHS function in the lhs package in R to generate the random sample of initial proportions for each compartment. This plot shows the initial trajectories of predicted cases for each study site. All other model compartments also converged after ~90 days.
